# Examining sex-differentiated genetic effects across neuropsychiatric and behavioral traits

**DOI:** 10.1101/2020.05.04.076042

**Authors:** Joanna Martin, Ekaterina A. Khramtsova, Slavina B. Goleva, Gabriëlla A M. Blokland, Michela Traglia, Raymond K. Walters, Christopher Hübel, Jonathan R I. Coleman, Gerome Breen, Anders D. Børglum, Ditte Demontis, Jakob Grove, Thomas Werge, Janita Bralten, Cynthia M. Bulik, Phil H. Lee, Carol A. Mathews, Roseann E. Peterson, Stacey J. Winham, Naomi Wray, Howard J. Edenberg, Wei Guo, Yin Yao, Benjamin M. Neale, Stephen V. Faraone, Tracey L. Petryshen, Lauren A. Weiss, Laramie E. Duncan, Sex Differences Cross-Disorder Analysis Group of the Psychiatric Genomics Consortium, Jill M. Goldstein, Jordan W. Smoller, Barbara E. Stranger, Lea K. Davis

## Abstract

**Background:** The origin of sex differences in prevalence and presentation of neuropsychiatric and behavioral traits is largely unknown. Given established genetic contributions and correlations across these traits, we tested for a sex-differentiated genetic architecture within and between traits.

**Methods:** Using genome-wide association study (GWAS) summary statistics for 20 neuropsychiatric and behavioral traits, we tested for differences in SNP-based heritability (h^2^) and genetic correlation (r_g_<1) between sexes. For each trait, we computed z-scores from sex-stratified GWAS regression coefficients and identified genes with sex-differentiated effects. We calculated Pearson correlation coefficients between z-scores for each trait pair, to assess whether specific pairs share variants with sex-differentiated effects. Finally, we tested for sex differences in between-trait genetic correlations.

**Results:** With current sample sizes (and power), we found no significant, consistent sex differences in SNP-based h^2^. Between-sex, within-trait genetic correlations were consistently high, although significantly less than 1 for educational attainment and risk-taking behavior. We identified genome-wide significant genes with sex-differentiated effects for eight traits. Several trait pairs shared sex-differentiated effects. The top 0.1% of genes with sex-differentiated effects across traits overlapped with neuron- and synapse-related gene sets. Most between-trait genetic correlation estimates were similar across sex, with several exceptions (e.g. educational attainment & risk-taking behavior).

**Conclusions:** Sex differences in the common autosomal genetic architecture of neuropsychiatric and behavioral phenotypes are small and polygenic, requiring large sample sizes. Genes with sex-differentiated effects are enriched for neuron-related gene sets. This work motivates further investigation of genetic, as well as environmental, influences on sex differences.

## Introduction

Despite widespread evidence of sex differences across human complex traits, including neuropsychiatric and behavioral phenotypes [1], the etiology of these differences remains poorly understood. Accumulating evidence suggests that sex differences in complex human phenotypes are likely to include a genetic component beyond that contributed by sex chromosomes and hormones [2–5]. Understanding the biological basis of sex differences in human disease, including neuropsychiatric disorders and traits, is critical for developing sex-informed diagnostics and therapeutics and realizing the promise of precision medicine [4]. Moreover, genetic variants with sex-differentiated effects across multiple traits may influence patterns of comorbidity for neuropsychiatric disorders and related behavioral traits, suggesting the need for cross-disorder genetic analyses to be evaluated in the context of sex-specific effects [6–11].

Neuropsychiatric and behavioral phenotypes are generally characterised by a complex and highly polygenic etiology [12]. Many of these traits share common variant genetic risks [13,14]. Specific genetic loci with pleiotropic effects are known to impact risk for multiple related neuropsychiatric and behavioral phenotypes [12]. However, it is not yet known whether these pleiotropic effects are consistent across females and males.

Recent studies have begun to investigate sex-differentiated genetic effects for a number of neuropsychiatric traits (see references in **Table 1**). Given evidence of phenotypic sex differences in prevalence and presentation, as well as genetic correlations across these traits [13], we set out to systematically test the hypothesis that neuropsychiatric and behavioral phenotypes with evidence of sex differences have a partially sex-differentiated autosomal genetic architecture that may be shared across traits. In this study, we have characterized the: (1) sex-dependent genetic architecture for a range of neuropsychiatric and behavioral traits, (2) degree of shared genetic architecture between males and females within each phenotype, and (3) sex-specific patterns of genetic effects shared across traits; see **Figure 1** for an overview of the analyses.

**Figure 1.**
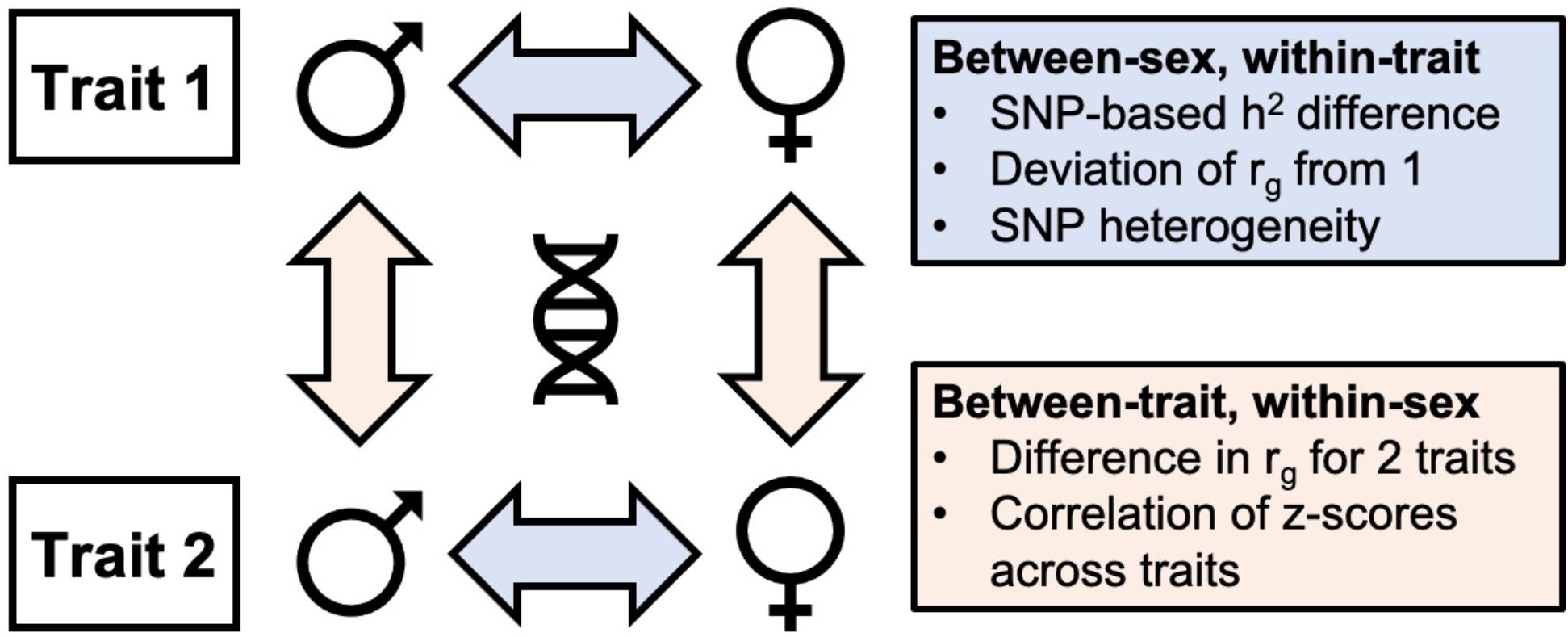
Schematic illustration of the key analyses used to investigate between-sex, within-trait and between-trait, within-sex differences.

**Table 1:** Summary of analyzed datasets of neuropsychiatric and behavioral traits

| Phenotype (full name) | Acronym | # of Female Cases | # of Female Controls | # of Male Cases | # of Male Controls | M:F case ratio | Sample type | Reference |
| --- | --- | --- | --- | --- | --- | --- | --- | --- |
| Attention-deficit hyperactivity disorder | ADHD | 4,945 | 16,246 | 14,154 | 17,948 | 2.86 | Clinical case-control | [35] |
| Alcohol dependence | ALCD | 2,504 | 6,033 | 5,932 | 9,412 | 2.37 | Clinical case-control | [42] |
| Anxiety disorders | ANX | 3,148 | 191,005 | 1,813 | 165,175 | 0.58 | General population (UK) | [43] |
| Autism spectrum disorder | ASD | 7,498 | 24,309 | 30,168 | 32,417 | 4.02 | Clinical case-control | [32,44] |
| Bipolar disorder | BD | 10,879 | 14,226 | 7,406 | 13,573 | 0.68 | Clinical case-control | [30] |
| Cannabis use (ever) | CUE | 17,244 | 71,742 | 17,414 | 50,737 | 1.01 | General population (UK) | N/A |
| Insomnia | INS | 19,521 | 39,846 | 12,863 | 40,776 | 0.66 | General population (UK) | [45] |
| Major depressive disorder | MDD | 10,711 | 11,745 | 5,021 | 11,226 | 0.47 | Clinical and population case-control | [30] |
| Major depressive disorder | N/A* | 13,492 | 180,661 | 7,156 | 159,832 | 0.53 | General population (UK) | [43] |
| Major depressive disorder recurrent | MDDR | 6,739 | 8,949 | 2,934 | 8,162 | 0.44 | Clinical case-control | [30] |
| Obsessive compulsive disorder | OCD | 1,525 | 4,307 | 1,249 | 2,789 | 0.82 | Clinical case-control | [46] |
| Post-traumatic stress disorder | PTSD | 968 | 2,457 | 585 | 4,025 | 0.60 | Clinical case-control | [47] |
| Risk-taking behavior | RTB | 32,285 | 143,678 | 51,392 | 100,984 | 1.59 | General population (UK) | [48] |
| Schizophrenia | SCZ | 9,854 | 16,785 | 18,366 | 17,122 | 1.86 | Clinical case-control | [30] |
| Smoking (current) | SMKC | 16,995 | 176,392 | 20,093 | 146,226 | 1.18 | General population (UK) | [43] |
| Smoking (previous) | SMKP | 62,305 | 131,082 | 65,245 | 101,074 | 1.05 | General population (UK) | [43] |
| Phenotype | Acronym | # of Females |  | # of Males |  | M:F ratio | Sample type | Reference |
| Alcohol use | ALCC | 59,088 |  | 53,088 |  | 0.90 | General population (UK) | [49] |
| Alcohol use | N/A* | 85,800 |  | 55,120 |  | 0.64 | General population | [50] |
| Age at first birth | AFB | 189,656 |  | 48,408 |  | 0.26 | General population | [51] |
| Educational attainment | EA | 182,286 |  | 146,631 |  | 0.80 | General population | [52] |
| Number of children ever born | NEB | 225,230 |  | 103,909 |  | 0.46 | General population | [51] |
| Neuroticism | NEU | 144,660 |  | 142,875 |  | 0.99 | General population (UK) | [33] |

\* These summary statistics were not used for analysis (see Supplemental Text for details).
PGC: Psychiatric Genomics Consortium; iPSYCH: The Lundbeck Foundation Initiative for Integrative Psychiatric Research.

## Methods & Materials

### Datasets

We collected sex-stratified GWAS meta-analysis summary statistics for 20 neuropsychiatric and behavioral traits (see **Table 1 & Supplemental Text**), chosen based on data availability. See **Table S1** for information about data availability. We used a broad definition of brain-based human complex traits, given the overwhelming evidence of shared genetic effects across such traits [13].

We used results from European-only meta-analyses to minimize any bias that may arise from ancestry differences, and because sufficiently large sex-stratified summary data of other ancestries are not currently available for the majority of these traits.

### Estimating sex-specific SNP-based heritability

For each trait, we calculated sex-specific observed scale SNP-based heritability (h^2^) using linkage disequilibrium (LD) score regression (LDSC) with pre-computed European ancestry LD scores (excluding SNPs in the HLA/MHC region; chr6:25-34M) [15]. For 11 binary traits, we also estimated SNP-based h^2^ on the liability scale, using sex-specific population prevalence rates from two sources, as described below. For comparison, we also used a second method, LDAK-SumHer [16], to estimate SNP-based h^2^, using the LDAK (LD-adjusted kinships) heritability model.

#### Sex-specific trait prevalence

We obtained sex-specific trait prevalence estimates from the USA and cumulative incidence rates from Denmark. The US-based estimates were derived from a hospital-based cohort of 752,436 patients who meet a medical home definition for Vanderbilt University Medical Center and whose de-identified electronic health record (EHR) is in the clinical research database [17]; see **Tables S2 & S3** for details. The medical home is a heuristic definition aimed at reducing the influence of missing data. Individuals meeting the medical home definition must have at least 5 ICD codes assigned on unique days over the span of at least 3 years. We used estimates based on adults (age >18 years), except for ADHD & ASD, which included pediatric patients. To estimate prevalence of each phenotype (**Supplementary Table 2**), we included individuals with at least one ICD code for each phenotype as the numerator, and ‘medical home’ hospital population as the denominator. However, hospital-based population prevalence estimates may be biased due to over-representation of individuals with more severe health-related conditions and higher levels of comorbidity. Additionally, these prevalence estimates may not generalize to populations outside of the USA. Therefore, we also used sex-specific cumulative incidence rates at age 50 years, based on individuals identified through inpatient and outpatient care in Denmark [18], as well as childhood-specific (age <18 years) estimates for the 2 neurodevelopmental disorders in our analyses (ADHD & ASD), based on a more recent Danish study, as the discovery GWAS for these traits were based on samples of mostly children [19]; see **Table S2**.

#### Statistical analysis of sex differences

For traits with non-zero SNP-based h^2^ estimates (i.e. where confidence intervals did not overlap with zero) in both sexes, we tested whether these sex-specific SNP-based h^2^ estimates were significantly different, by calculating z-scores using Equation 1 (below), and obtaining corresponding p-values from a normal distribution. We corrected for multiple tests using a Bonferroni correction for N=12 independent tests (N=5 continuous traits and N=7 binary traits that were non-zero in both sexes and were converted to the liability scale; p-value threshold = 0.0042).

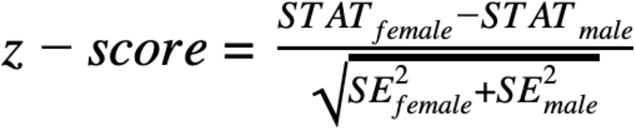

*Equation 1, where STAT can be any statistic for which we want to assess the difference between the sexes, including SNP-based h^2^, r_g_, and GWAS betas; SE is the standard error for the statistic. This test will be well calibrated as long as STAT/SE is normally distributed and the test statistics are independent between sexes, and will be conservative if the statistics are positively correlated.*

### Estimating genetic correlations

We used LDSC to estimate genetic correlations (r_g_): 1) between sexes, within each trait and 2) between each trait pair, within sex (see **Figure 1**). For between-sex, within-trait correlations, we tested the null hypothesis that r_g_ was significantly lower than 1 using a one-tailed test compared to a normal distribution (z=(1-r_g_)/SE). We applied a Bonferroni correction to account for multiple tests (p<0.0031 based on 16 traits). Next, we tested whether the between-trait r_g_ estimates were different for males (r_gM_) and females (r_gF_), by using a z-score approximation based on block jackknife to estimate the standard error of r_gM_-r_gF_ in LDSC. As with other LDSC analyses, this approach is robust to sample overlap. We applied a false discovery rate (FDR) correction to the results to account for multiple tests, given the large number of non-independent genetic correlations across phenotypes, which would make a Bonferroni correction overly conservative. Genetic correlations were visualized using the Python package Networkx [20] and matplotlib [21].

### Between-sex, within-trait genetic heterogeneity

For each SNP in the sex-stratified GWAS of each trait, we assessed between-sex, within-trait heterogeneity using z-scores (which are correlated with Cochran’s Q statistic but provide directionality of the effect) as in Equation 1. This test quantifies the difference in SNP association effect size between the sexes, similar to, although not the same as, an interaction test [22]. Given that only summary statistics from sex-stratified GWAS were available, the analysis of sex differentiated genetic effects was limited to the z-score approach.

### Sharing of variants with sex-differentiated effects across traits

To assess which traits share variants with sex-differentiated effects (i.e. variants at the extreme ends of the z-score distribution), we assessed the Pearson correlation coefficient between z-scores (i.e. the differences of betas from male-only and female-only GWAS) for pairs of traits. Given that there are many non-independent observations, due to SNPs in LD, we used a block jackknife approach to estimate the significance of the Pearson correlation [23,24]. SNPs were assigned to one of 1000 contiguous blocks based on their genomic position. For each pair of traits, Pearson correlation was calculated on the full set of z-scores and then recalculated after each block was removed, thus estimating the jackknife error and p-values.

### Gene-based analysis, functional mapping and gene-set enrichment analysis of genes with sex-differentiated effects

We used the Functional Mapping and Annotation of Genome-Wide Association Studies (FUMA) SNP2GENE web tool [25], to perform gene-based analysis and positional mapping of variants to genes. Z-scores computed for each trait were used as an input. For the gene-based analysis implemented via generalized-gene set analysis of GWAS data (MAGMA) [26] in FUMA, we used the default setting in which SNPs are mapped to genes if they fall within a window spanning 10kb before the start and after the end positions for the gene. In this analysis, the mean of the chi^2^ statistic for the SNPs in a gene is calculated and the p-value is obtained from a known approximation of the sampling distribution [27,28]. The genome-wide significance threshold is defined as 0.05 / number of genes to which the SNPs are mapped.

After mapping SNPs to genes, we filtered the top 0.1% (with the lowest -log10(p-value) from the gene-based MAGMA analysis) as genes with sex-differentiated effects for each trait and combined these sets across phenotypes. We selected 0.1% as the cut-off in order to test a set of genes with the greatest sex difference in effects, that did not exceed ~2000 genes, which is an input cut-off for the Gene Set Enrichment Analysis (GSEA) tool. We also tested 0.5% as a sensitivity check, which results in similar genes sets, but picks up a broader set of genes given a larger list of genes. Next we computed a gene set overlap analysis using GSEA (https://software.broadinstitute.org/gsea/index.jsp) on the combined set of genes with sex-differentiated effects with collection C5 (GO biological process, GO cellular component, and GO molecular functions) from MSigDB to investigate which gene sets may contribute to phenotypic differences observed for the neuropsychiatric and behavioral traits.

## Results

### Sex-stratified SNP-based h^2^ estimates and assessment of sex differences

Sex-specific SNP-based h^2^ estimates using LDSC are presented in **Figure 2**, with details provided in **Table S4**. Several traits (post-traumatic stress disorder (PTSD) & recurrent major depressive disorder (recurrent MDD) in males and autism spectrum disorder (ASD) & alcohol dependence in females) did not have sufficient power (or may have been affected by a high degree of heterogeneity) and we did not detect a polygenic signal using LDSC and therefore sex differences could not be assessed. Thus, although we report sex difference estimates for all traits in **Table S4**, these cannot be reliably interpreted for ASD, recurrent MDD, alcohol dependence, and PTSD, since one of the sexes exhibited a near zero or negative SNP-based h^2^ estimate. The liability scale SNP-based h^2^ estimates using population prevalence from the USA and cumulative incidence from Denmark were highly correlated (r^2^=0.97, p=5.1×10^-10^); see **Figure S1**. Age at first birth of child was the only trait with a significant sex difference (after multiple testing correction; p<0.0042) in SNP-based h^2^ estimates (females: SNP-based h^2^=0.052, SE=0.004; males: SNP-based h^2^=0.113, SE=0.010); (z-score=-5.81, p=6.43×10^-9^).

**Figure 2.**
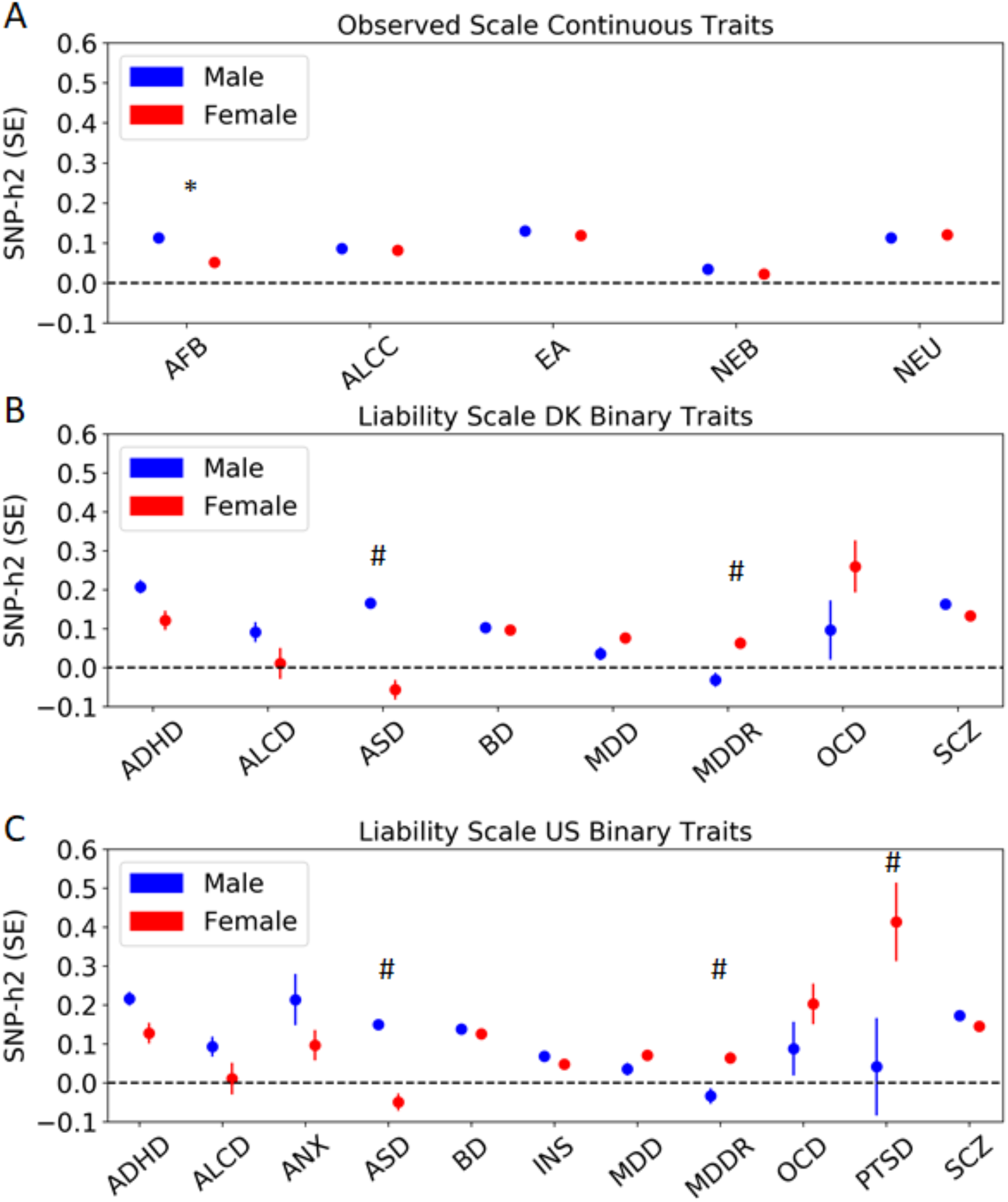
Estimates of sex stratified SNP-based heritability (h^2^) on (A) the observed scale for continuous traits, and the liability scale, using population prevalence based on (B) DK (Denmark) and C) the USA. Estimates were obtained from LDSC. Points represent the estimated SNP-based h^2^ in males (blue) and females (red), while bars represent standard errors (SE) of the SNP-based h^2^ estimates. Significant sex difference in heritability is denoted with an asterisk, as follows: * p<0.0042 (adjusted p-value threshold corrected for multiple testing using Bonferroni). # denotes traits for which significance in difference is not interpretable due to negative or non-significant from zero SNP-based h^2^ value for one of the measurements. Phenotype abbreviations are as follows: ADHD: Attention-deficit hyperactivity disorder; AFB: Age at first birth; ALCC: Alcohol use; ALCD: Alcohol dependence; ANX: Anxiety disorders; ASD: Autism spectrum disorders; BD: Bipolar disorder; EA: Educational attainment; INS: Insomnia; MDD: Major depressive disorder; MDDR: Major depressive disorder recurrent; NEB: Number of children ever born; NEU: Neuroticism; OCD: Obsessive compulsive disorder; PTSD: Post-traumatic stress disorder; SCZ: Schizophrenia.

Observed scale SNP-based h^2^ estimates based on LDAK-SumHer were somewhat higher than those obtained in LDSC and moderately correlated with them (r^2^=0.53, p=4.4×10^-4^ for all traits, r^2^=0.73, p=4.1×10^-7^ excluding traits for which SNP-based h^2^ could not be reliably estimated in LDSC, i.e. PTSD & recurrent MDD in males and ASD & alcohol dependence in females); see **Table S5** and **Figures S1 & S2** for details. Higher estimates from the LDAK model relative to the LDSC model have been previously observed for a variety of traits [16,29]. In contrast to LDSC results, age at first birth did not show a significant sex difference after multiple testing correction (z-score=1.94, p=0.052), with an effect in the opposite direction to that observed using LDSC. Using LDAK, the liability scale (adjusted based on each population) SNP-based h^2^ estimates differed by sex for the following traits: recurrent MDD (US: z-score=-4.57, p=4.81×10^-6^; DK: z-score=-4.36, p=1.33×10^-5^), ASD (US: z-score=2.94, p=0.0033; DK: z-score=3.28, p=0.0011), and schizophrenia (DK: z-score=-3.16, p=0.0016). These results were not observed using LDSC, and indeed SNP-based h^2^ could not be estimated reliably in LDSC for ASD in females or recurrent MDD in males. The biggest discrepancies between estimates obtained from LDSC and LDAK were for the traits with the smallest sample sizes (see **Figure S3**).

The SNP-based h^2^ results for ADHD and ASD were similar albeit somewhat higher for both LDSC and LDAK when using estimates based on a Danish child-specific study [19], compared to using prevalence estimates from the whole Danish population [18]; see **Tables S4 and S5**.

### Between-sex, within-trait genetic correlation analysis

We quantified the genetic correlation between males and females for each trait (excluding the four traits where SNP-based h^2^ could not be estimated in both sexes); see **Figure 3 and Table S6**. We found moderate to high genetic correlations for all traits (r_g_ = 0.68 – 1.21); these all differed significantly from zero and we also detected a significant difference from 1 for risk-taking behavior: r_g_(se)=0.81 (0.04) and educational attainment: r_g_(se)=0.92(0.02), after correcting for multiple tests (p<0.0031), suggesting a modest degree of common variant heterogeneity in males and females for these phenotypes.

**Figure 3.**
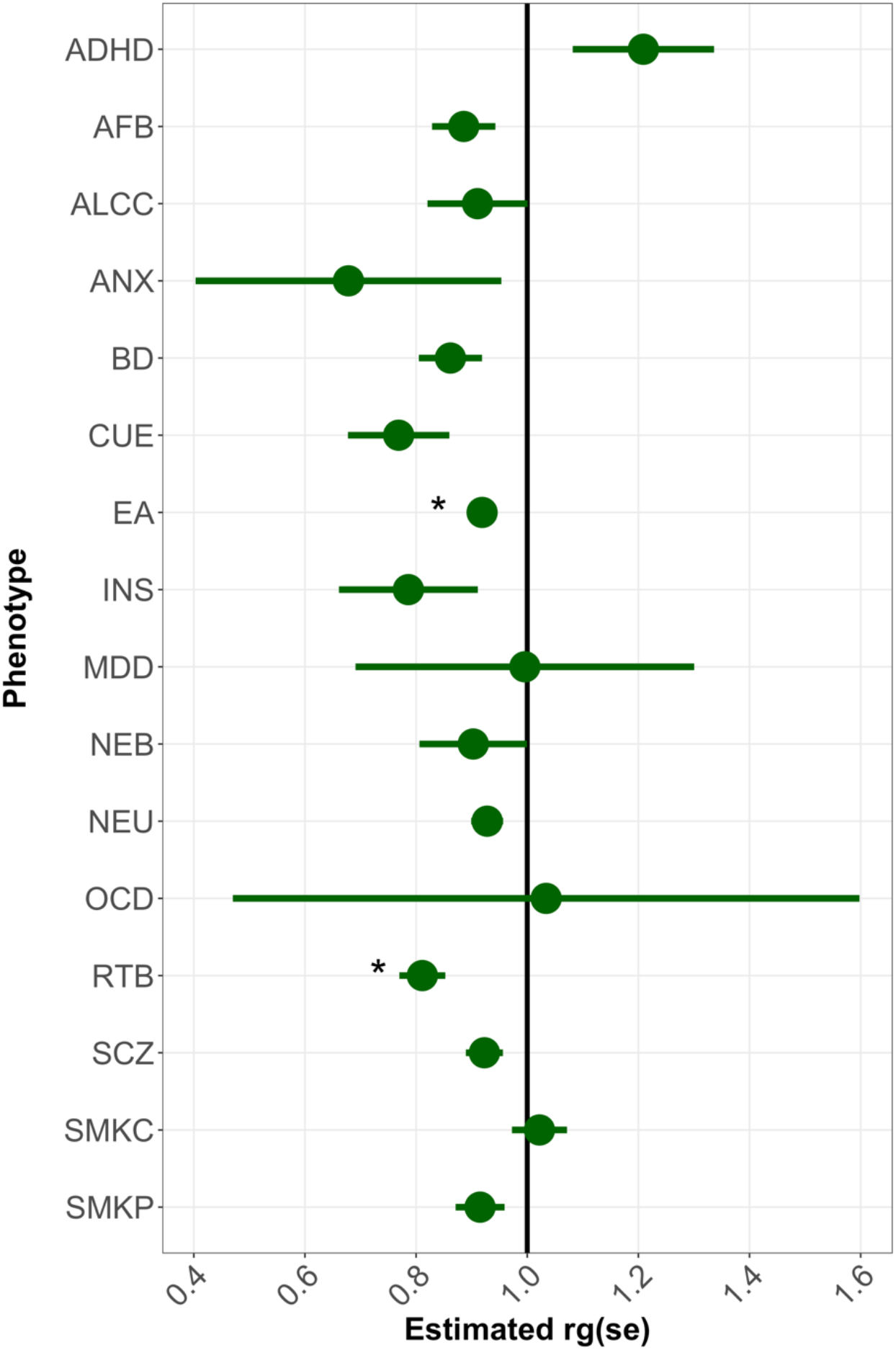
Within-trait, between-sex genetic correlation (r_g_) estimates using LDSC. Points represent the estimated r_g_ and bars represent standard errors (SE) of the r_g_ estimates. Significant deviation from 1 is denoted with an asterisk, as follows: * p<0.0031 (adjusted p-value threshold corrected for multiple testing using Bonferroni). Phenotype abbreviations are as follows: ADHD: Attention-deficit hyperactivity disorder; AFB: Age at first birth; ALCC: Alcohol use; ANX: Anxiety disorders; BD: Bipolar disorder; CUE: Cannabis use (ever); EA: Educational attainment; INS: Insomnia; MDD: Major depressive disorder; NEB: Number of children ever born; NEU: Neuroticism; OCD: Obsessive compulsive disorder; RTB: Risk-taking behavior; SCZ: Schizophrenia; SMKC: Smoking (current); SMKP: Smoking (previous).

### Between-sex, within-trait heterogeneity across variants

To assess sex differences in genetic effects of common variants, for each trait we computed z-scores and corresponding p-values for each SNP, using Equation 1. **Figure S4** shows the quantile-quantile (QQ) plots of the z-score p-values for all traits. While the difference in the beta estimates between males and females did not reach genome-wide statistical significance (5×10^-8^) for any given SNP, we did observe deviation from the expected null distribution (**Figure S4**) for ADHD, lifetime cannabis use, MDD, number of children born, and schizophrenia. **Figure 4A** shows a Miami plot for female-only (top) and male-only (bottom) lifetime cannabis use GWAS, where we observed several associations that are stronger in females (e.g. on chromosomes 3, 6, 16 and 18). Since cohorts for lifetime cannabis use are of very similar size the power to detect association in both sexes is similar.

A gene-based analysis in FUMA revealed several traits with genome-wide significant genes with sex-differentiated effects. Gene-based analysis Manhattan plots are shown in **Figure S5**. Traits with significant gene associations include alcohol consumption (gene: *PELI2),* alcohol dependence *(ADAM23),* anxiety *(PRKCH* and *KLHDC4),* lifetime cannabis use *(MYOF),* number of children born *(GLB1L2),* neuroticism *(EXTL2),* risk-taking behavior *(HFE2* and *AGO2*), and schizophrenia *(SLTM).* Interestingly, *SLTM*gene, which is highly expressed in cerebellum (GTEx Portal, www.gtexportal.org), was also identified in a gene-based gene-by-sex interaction for schizophrenia and across schizophrenia, bipolar disorder, and MDD disorders [30].

### Shared sexually-differentiated effects across traits

To assess whether specific pairs of traits share sex-differentiated effects, for each pair of traits we calculated the Pearson correlation coefficient between each trait’s SNP z-scores for sex-differentiated effects. **Figure 4B** shows a matrix of Pearson correlation coefficients for pairs of traits. We find small to moderate, but significant, correlations of z-scores for several traits. Interestingly, we find that the correlation of z-scores between MDD and recurrent MDD is high, but not equal to 1 (Pearson correlation coefficient = 0.77, p<0.001), indicating that there are both shared and trait-specific variants with sex-differentiated effects for these two definitions of MDD. Furthermore, we find cross-trait sharing of sex-dependent genetic effects between ASD and ADHD and also bipolar disorder and schizophrenia, to name a few examples.

**Figure 4.**
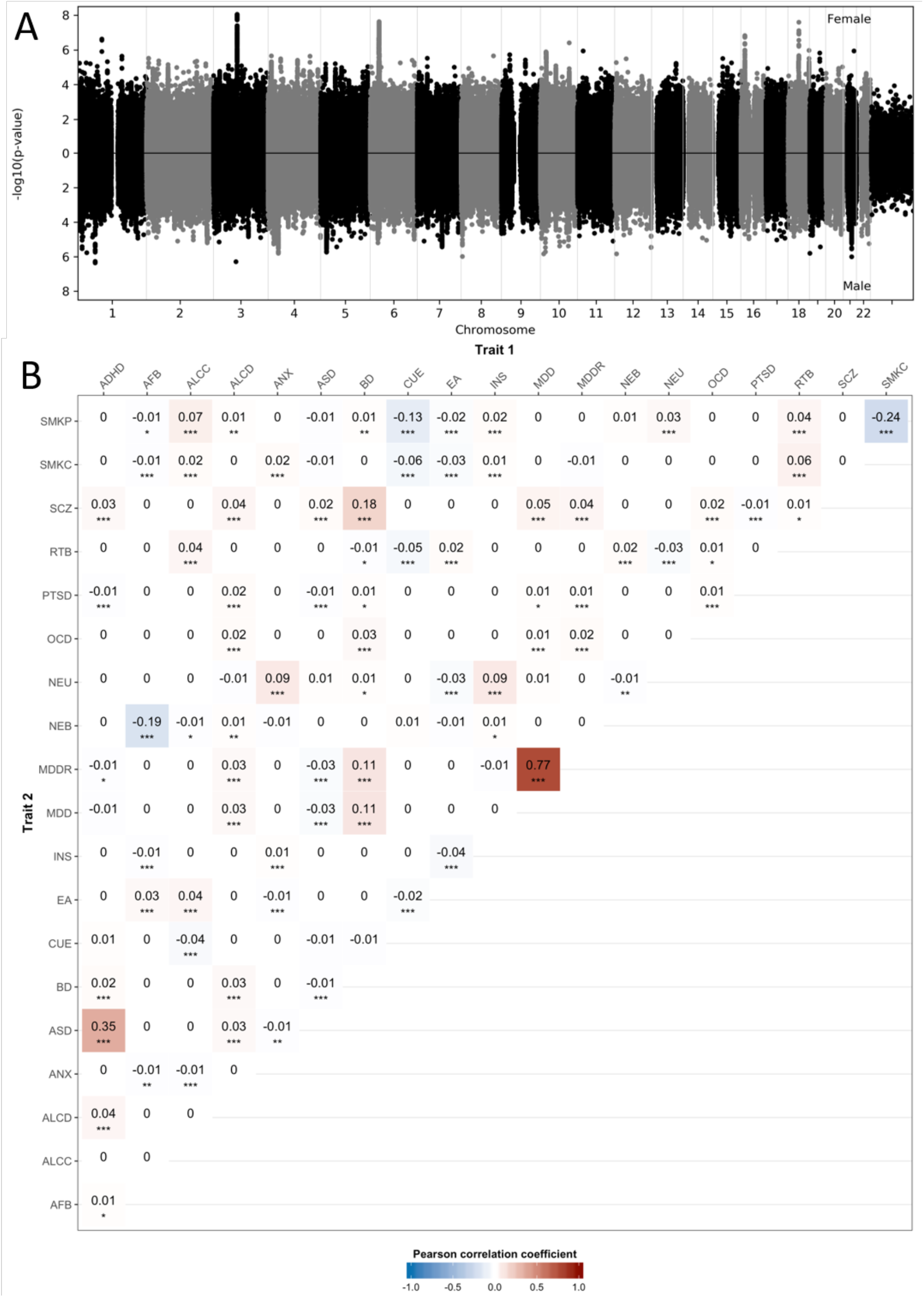
Sharing of variants with sexually-differentiated effects between-traits. (A) Miami plot for female-only (top) and male-only (bottom) GWAS for cannabis use (ever); female cases: N=17,244; male cases: N=17,414. For each SNP, we computed z-scores using Equation 1. (B) Matrix of Pearson correlation coefficients for pairs of traits. We performed Pearson correlation of z-scores and a block jackknife approach to estimate the significance of the correlation for all pairs of traits. Asterisks indicate the estimated significance of the coefficients, as follows: * p<0.05, **p<0.01, ***p<0.001. Color coding represents positive (red) or negative (blue) correlation. Phenotype abbreviations are as follows: ADHD: Attention-deficit hyperactivity disorder; AFB: Age at first birth; ALCC: Alcohol use; ALCD: Alcohol dependence; ANX: Anxiety disorders; ASD: Autism spectrum disorders; BD: Bipolar disorder; CUE: Cannabis use (ever); EA: Educational attainment; INS: Insomnia; MDD: Major depressive disorder; MDDR: Major depressive disorder recurrent; NEB: Number of children ever born; NEU: Neuroticism; OCD: Obsessive compulsive disorder; PTSD: Post-traumatic stress disorder; RTB: Risk-taking behavior; SCZ: Schizophrenia; SMKC: Smoking (current); SMKP: Smoking (previous).

### Gene set overlap analysis of genes with sex-differentiated effects across traits

To investigate the biological function of the genes harboring SNPs with sex-differentiated genetic effects, we selected the top 0.1% of genes from each trait (**Table S7**), resulting in 349 genes that were mapped for GSEA. The gene sets overlapping genes with sex-differentiated effects (FDR q-value < 0.01) are listed in **Table S8.** Interestingly, the gene sets significantly enriched for genes with sex-differentiated effects (FDR q-value < 0.01) included neurogenesis, regulation of neuron projection development, signaling receptor binding, regulation of neuron differentiation, neuron differentiation, and neuron development, and synapse maturation gene sets.

### Between-trait, within-sex genetic correlation analysis

The genetic correlation results are presented as network plots (**Figure 5**) and heatmaps (**Figure S6**). The overall pattern of between-trait genetic correlations was similar in males and females (**Figure 5B, C**). We detected several significant sex differences in between-trait genetic correlations; see **Table 2** and **Figure 5A** for top results and **Table S9** for full details. The genetic correlation (r_g_) between educational attainment and risk-taking behavior was positive in females but negative in males, while for lifetime cannabis use and neuroticism, the correlation was negative in females but positive in males. The magnitude of r_g_ was significantly greater in females than males for a number of traits (e.g. risk-taking behavior & schizophrenia) and significantly smaller in females than males for several trait pairs (e.g. number of children born & risk-taking behavior). However, for a number of these trait pairs, the estimated r_g_ in one sex did not significantly differ from zero (see **Table S9**), suggesting that either there was no significant genetic correlation between a given pair of traits in one sex or the power to estimate this effect was too low.

**Figure 5.**
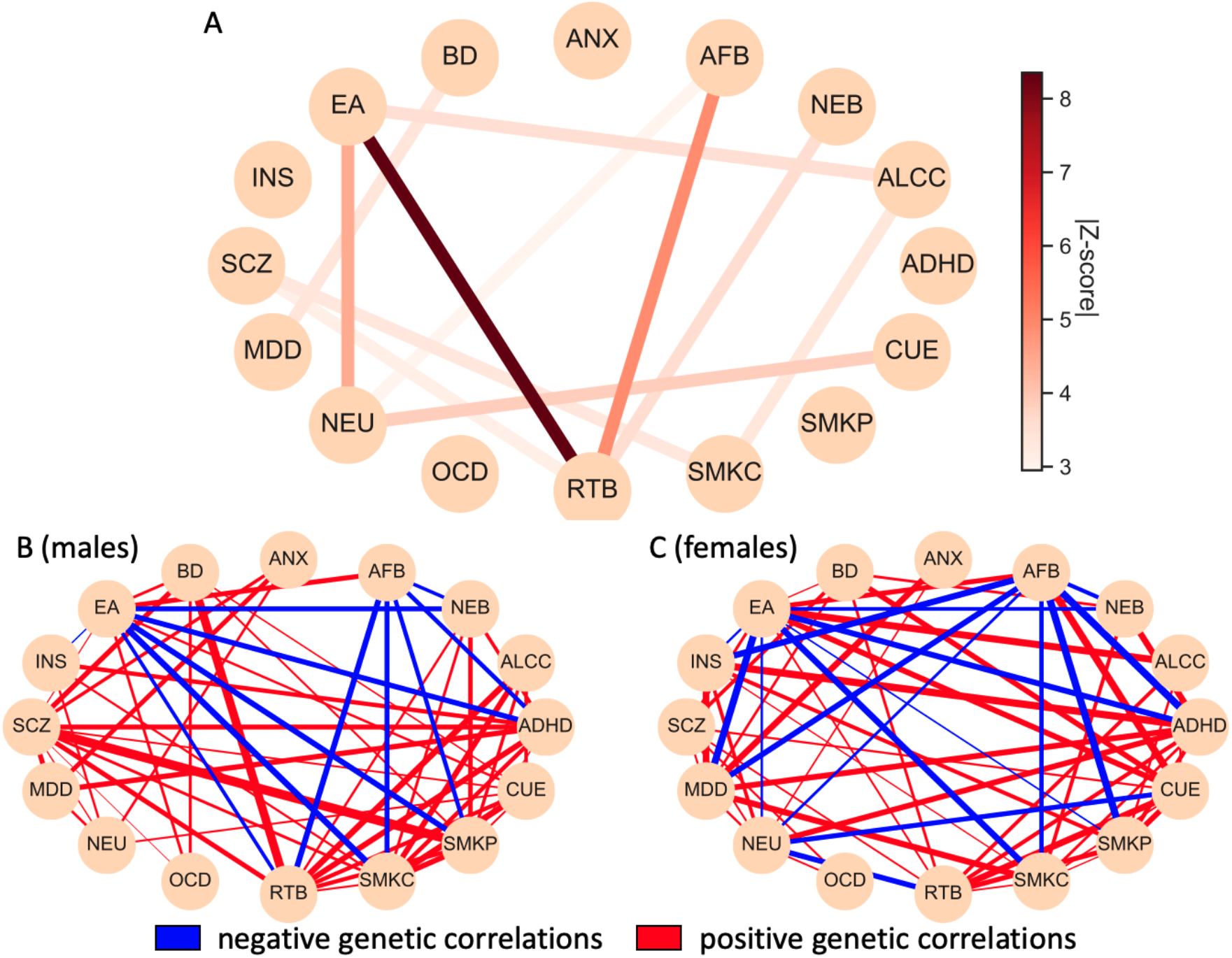
(A) A network plot showing between-trait genetic correlations with a significant sex difference as computed by a z-score. The edge color represents the absolute value of the z-score for the difference in genetic correlation between the same two phenotypes in females vs. males. Only pairs of traits with an FDR corrected q<0.05 sex difference are shown. (B & C) Between-trait, within-sex genetic correlation analysis. Network plots for genetic correlation estimates (r_g_) for pairs of traits in (B) males and (C) females, where each node represents a trait, and the edge represents positive (red) or negative (blue) genetic correlation. The thickness of the edge represents -log10(q-value) of correlation significance. Only genetic correlations with FDR corrected q<0.05 are shown. Phenotype abbreviations are as follows: ADHD: Attention-deficit hyperactivity disorder; AFB: Age at first birth; ALCC: Alcohol use; ANX: Anxiety disorders; ASD: Autism spectrum disorders; BD: Bipolar disorder; CUE: Cannabis use (ever); EA: Educational attainment; INS: Insomnia; MDD: Major depressive disorder; NEB: Number of children ever born; NEU: Neuroticism; OCD: Obsessive compulsive disorder; RTB: Risk-taking behavior; SCZ: Schizophrenia; SMKC: Smoking (current); SMKP: Smoking (previous).

**Table 2:** Top results of sex differences in cross-trait genetic correlation estimates

| Trait 1 | Trait 2 | Females |  |  | Males |  |  | Sex difference |  |
| --- | --- | --- | --- | --- | --- | --- | --- | --- | --- |
| | | $r_g$ | SE | q-value<sub>R</sub> | $r_g$ | SE | q-value<sub>R</sub> | z-score | q-value |
| EA | RTB | 0.187 | 0.033 | $6.38 \times 10^{-8}$ | -0.144 | 0.033 | $4.29 \times 10^{-5}$ | -8.353 | $7.98 \times 10^{-15}$ |
| AFB | RTB | -0.035 | 0.046 | 0.52 | -0.344 | 0.054 | $1.23 \times 10^{-9}$ | -4.906 | $5.58 \times 10^{-5}$ |
| EA | NEU | -0.22 | 0.029 | $1.72 \times 10^{-13}$ | -0.064 | 0.029 | 0.051 | 4.421 | $3.94 \times 10^{-4}$ |
| CUE | NEU | -0.142 | 0.055 | 0.022 | 0.124 | 0.054 | 0.044 | 3.866 | $3.32 \times 10^{-3}$ |
| NEB | RTB | 0.116 | 0.063 | 0.12 | 0.413 | 0.074 | $1.43 \times 10^{-7}$ | 3.582 | $8.19 \times 10^{-3}$ |
| ALCC | EA | 0.276 | 0.047 | $2.52 \times 10^{-8}$ | 0.043 | 0.049 | 0.47 | -3.53 | $8.30 \times 10^{-3}$ |
| SCZ | SMKC | 0.034 | 0.045 | 0.52 | 0.214 | 0.046 | $1.54 \times 10^{-5}$ | 3.301 | 0.013 |
| ALCC | SMKC | 0.013 | 0.058 | 0.86 | 0.292 | 0.069 | $8.97 \times 10^{-5}$ | 3.326 | 0.013 |
| BD | MDD | 0.565 | 0.079 | $4.95 \times 10^{-12}$ | 0.057 | 0.142 | 0.74 | -3.367 | 0.013 |
| RTB | SCZ | 0.326 | 0.043 | $3.13 \times 10^{-13}$ | 0.157 | 0.038 | $1.07 \times 10^{-4}$ | -3.088 | 0.024 |
| AFB | NEU | -0.173 | 0.037 | $1.44 \times 10^{-5}$ | -0.028 | 0.048 | 0.63 | 2.95 | 0.035 |

Z-scores were calculated using Equation 1. AFB: Age at first birth; ALCC: Alcohol use; BD: Bipolar disorder; CUE: Cannabis use (ever); EA: Educational attainment; MDD: Major depressive disorder; NEB: Number of children ever born; NEU: Neuroticism; RTB: Risk-taking behavior; SCZ: Schizophrenia; SMKC: Smoking (current).

## Discussion

We investigated sex differences in the genetic architecture of 20 neuropsychiatric and behavioral traits, using sex-stratified autosomal GWAS summary statistics. We used three complementary approaches, including estimation of SNP-based heritability, genetic correlation, and heterogeneity analyses, to evaluate sex differences within traits and across pairs of traits. As expected, most common autosomal genetic effects are shared across sexes. However, a number of notable sex differences were detected.

For a large number of traits and cross-trait pairs, we detected no consistent evidence of sex differences in SNP-based heritability and the genetic correlations between males and females were moderate-to-high (mostly r_g_>0.8). The phenotypes that showed sex differences were among those with the largest available sample sizes, indicating that large sample sizes make the detection of sex differences more likely and consequently, the lack of significant sex differences for a given phenotype may be due to small sample sizes. For example, a recent analysis of schizophrenia, bipolar disorder and MDD with a larger sample size has revealed significant associations for schizophrenia and MDD [30]. We found that some pairs of genetically correlated traits also share sex-differentiated associations (e.g. ASD and ADHD; bipolar disorder and schizophrenia). Taken together, these findings suggest that sex differences in the genetic architecture of neuropsychiatric and behavioral traits exist, but are small and polygenic. They further support the hypothesis that SNPs with sex-differentiated genetic effects for one trait are also likely to exhibit sex-differentiated effects in phenotypically associated traits [31,32]. Moreover, we found that the genes with the most sex-differentiated effects across all traits are enriched for neuron- and synapse-related gene functions.

For two specific traits with well-powered GWAS (educational attainment and risk-taking behavior), several interesting results emerged. Both educational attainment and risk-taking behavior demonstrated similar SNP-based h^2^ in males and females, indicating that there was no appreciable difference in the overall burden of genetic factors accounting for phenotypic variance in each sex. Also, neither trait demonstrated an excess of variants with sex-differentiated effects, showing that (at current sample sizes) there were few detectable sex differences in SNP effects for either trait. However, while the genetic correlation between males and females was high (educational attainment: 0.92(0.02), as previously reported [33]; risk-taking behavior: 0.81(0.04)), it was significantly less than 1 for both traits. Moreover, these two traits were positively genetically correlated with each other (r_g_=0.19) in females but negatively correlated in males (r_g_=-0.14). These results may be explained by circumstances in which there exists a large number of SNPs with very small sex-differentiated effects, which we remain underpowered to detect at individual loci, but can observe in analyses of cumulative sex differences. An alternative possibility is that there are sex differences in ascertainment and measurement of these two phenotypes (e.g. males and females interpreting the question about being a risk-taker differently), thus resulting in analysis of slightly different traits in males and females. In general, ascertainment effects (e.g. potential recruitment and participation biases) and measurement issues (e.g. phenotyping biases) should be carefully considered in future genetic studies of sex differences. Many of the GWAS of behavioral traits are based on data from UK Biobank (which is an older sample of relatively healthier and wealthier individuals compared to the general UK population) [34], whereas the case-control neuropsychiatric traits are frequently ascertained from clinical populations.

These observations have important implications for the future of sex differences research. Although the majority of genetic effects for neuropsychiatric and behavioral traits are similar for males and females, sex-differentiated genetic effects can be identified. The full characterization of these effects will require larger sample sizes than for detection of main effects because of reduced statistical power in assessing the interaction between sex and genotype. We expect that as sample sizes increase, sex differences will emerge but will be small in magnitude, reflecting the polygenic architecture of the phenotypes and the sex differences. Furthermore, the large sex differences in prevalence of psychiatric disorders are unlikely to be explained entirely by sex differences in common genetic associations or global burden of autosomal genetic factors. Additional studies investigating the interaction between cumulative genetic effects (including non-autosomal and rare variation) and the environment (e.g. hormonal and social factors) will be needed to understand the origins of these differences.

### Limitations and Considerations

We focused on neuropsychiatric and behavioral traits with available sex-stratified GWAS summary statistics. The comprised cohorts were of European ancestry and due to limited data we were unable to assess and compare results across ancestry. Furthermore, lack of access to genotype-level data restricted our analyses to methods developed for summary statistics. We also note that our analyses can be impacted by the number of cases and controls and the ratio of females to males in each cohort (e.g. [35,36]). Indeed, estimation of SNP-based h^2^ relies on several important assumptions (e.g. regarding the underlying genetic architecture and number of causal variants per LD block) [15,16] and can be influenced by many factors (e.g. sex-specific population prevalences, gender-dependent ascertainment methods for cases and controls, different sample sizes in males and females) [37–39]. Accurate estimation of sex-specific population prevalences is complex given that there could be sex differences in referral, with under-diagnosis in one sex (e.g. as is the case for ADHD [40]). To account for the difficulties in estimation of SNP-based h^2^, we used two different methods (LDSC & LDAK) and prevalence estimates from two different populations (Denmark & USA). Estimates based on the two different population prevalence estimates were highly correlated, but there were substantial differences in estimation based on either LDSC or LDAK, likely due to the different model assumptions related to genetic architecture; the biggest discrepancies were for the traits with the smallest sample sizes (see **Figure S5**); the true SNP-based h^2^ estimate is likely to fall in between these estimates. Furthermore, it is likely that some of the GWAS summary statistics may have included data from ‘super-screened’ and unscreened controls, which may have biased upwards the genetic correlation estimates [41].

The most direct method to identify SNPs with sex-dependent effects is to perform a genotype-by-sex interaction test. However, this requires individual-level genotype data. A sex-stratified analysis followed by a difference test, such as the z-score used here, is equivalent to a genotype-by-sex interaction test when there is no interaction between covariates (e.g. PCs, age) and the strata (e.g. male and female), and the trait variances are equivalent in the two strata [22]. If those assumptions hold, then our stratified analyses will be conservative. Conversely, if those assumptions are violated then our stratified analysis will be robust to those covariate interactions and differences in residual variances when evaluating whether the common variant effects are heterogeneous across sex. Subsequent testing in larger cohorts may illuminate whether these assumptions are violated and their impact on the interpretation of variants with sex-differentiated effects.

Another important limitation of our study is that we only assessed the genetic effects on the autosomes. The sex chromosomes are very frequently excluded from GWAS, due to special consideration required for quality control and analyses, with many methods not allowing for the inclusion of sex chromosomes, and summary statistics were not available for the present analyses.

## Conclusion

Through within- and between-trait analyses, we find evidence of sex-dependent autosomal effects for several neuropsychiatric and behavioral phenotypes among European ancestry cohorts. However, the effects are small and polygenic and therefore larger samples are needed for identifying such effects and understanding their functional contribution to complex traits. Furthermore, studies of sex differences taking into account non-autosomal and rare genetic variants, as well as environmental and hormonal influences, including ethnic and cultural differences are also needed.

## Acknowledgements

JM was supported by the Wellcome Trust (grant 106047) and a Sêr Cymru II COFUND Fellowship from the Welsh Government.

RW was supported by the NIH (5U01MH109539) and the Stanley Center for Psychiatric Research.

LKD was supported in part by NIH (R01NS102371-01A1, R01MH113362, U01HG009086, R01MH118223, RM1HG009034).

JRIC and GB were supported in part by the English National Institute for Health Research (NIHR) as part of the Maudsley Biomedical Research Centre (BRC). This study represents independent research partly funded by the NIHR BRC at South London and Maudsley NHS Foundation Trust and King’s College London. The views expressed are those of the author(s) and not necessarily those of the NHS, the NIHR or the Department of Health and Social Care. High performance computing facilities were funded with capital equipment grants from the GSTT Charity (TR130505) and Maudsley Charity (980).

The iPSYCH team was supported by grants from the Lundbeck Foundation (R165-2013-15320, R102-A9118, R155-2014-1724 and R248-2017-2003) and the universities and university hospitals of Aarhus and Copenhagen. The Danish National Biobank resource was supported by the Novo Nordisk Foundation. Data handling and analysis on the GenomeDK HPC facility was supported by NIMH (1U01MH109514-01 to ADB). High-performance computer capacity for handling and statistical analysis of iPSYCH data on the GenomeDK HPC facility was provided by the Center for Genomics and Personalized Medicine and the Centre for Integrative Sequencing, iSEQ, Aarhus University, Denmark (grant to ADB). ADB also received funding from the European Union H2020 Programme (H2020/2014-2020) under grant agreement No. 667302 (CoCA).

PL is supported by NIH (R00MH101367, R01MH119243).

SVF is supported by the European Union’s Seventh Framework Programme for research, technological development and demonstration under grant agreement no 602805, the European Union’s Horizon 2020 research and innovation programme under grant agreements No 667302 & 728018 and NIMH grants 5R01MH101519 and U01 MH109536-01

REP is supported by NIMH K01MH113848 and The Brain & Behavior Research Foundation NARSAD grant 28632 P&S Fund.

HE was supported by the NIH (grant: MH109532).

JB was supported by funding from the European Community’s Horizon 2020 research and innovation programme under grant agreement no. 847879 (PRIME).

CAM is supported by NINDS grants R01 NS102371 and R01 NS105746.

LW & MT were supported by grant R01MH114924.

This work utilized the computational resources of the Dutch national e-infrastructure with the support of SURF Cooperative (https://userinfo.surfsara.nl/).

With thanks to Dr Helena Gaspar for providing a python script that was used to estimate differences in LDSC r_g_ across sex, and to Donald Hucks for computing population prevalence by sex for the BioVU population.

The original GWAS data were supported by National Institute of Mental Health Psychiatric Genomics Consortium grants: **U01 MH109528**; U01 MH109539; U01 MH109536; U01 MH109501; **U01 MH109514**; U01 MH109499; U01 MH109532.

We would also like to thank the following consortia & groups that contributed data:

- Attention Deficit Hyperactivity Disorder Working Group of the Psychiatric Genomics Consortium (PGC) and the Lundbeck Foundation Initiative for Integrative Psychiatric Research (iPSYCH)
- Autism Spectrum Disorder Working Group of the PGC & iPSYCH
- Bipolar Disorder Working Group of the PGC
- Major Depressive Disorder Working Group of the PGC
- Obsessive Compulsive Disorder Working Group of the PGC
- Post-traumatic Stress Disorder Working Group of the PGC
- Schizophrenia Working Group of the PGC
- Substance Use Disorders Working Group of the PGC
- UK Biobank GWAS results generated by the Neale Lab (http://www.nealelab.is/uk-biobank/)

## Author contributions

<u>Study conception, design, analyses & writing</u>: Joanna Martin and Ekaterina Khramtsova.

<u>Analyses & writing</u>: Slavina Goleva and Gabriëlla Blokland.

<u>Analyses & editing</u>: Michela Traglia, Raymond Walters, Christopher Hübel, Jonathan Coleman, and Gerome Breen.

<u>Editorial group</u>: Anders Børglum, Ditte Demontis, Jakob Grove, Thomas Werge, Janita Bralten, Cynthia Bulik, Phil Lee, Carol A. Mathews, Roseann E. Peterson, Stacey Winham, Naomi Wray, Howard Edenberg, Wei Guo, Yin Yao, Benjamin Neale, Stephen V. Faraone, Tracey Petryshen, Lauren Weiss, and Laramie Duncan

<u>Writing</u>: Jill Goldstein and Jordan Smoller.

<u>Writing and analytic supervision</u>: Barbara Stranger & Lea Davis.

