## Supplemental Text & Figures for "Examining sex-differentiated genetic effects across neuropsychiatric and behavioral traits"

### Supplemental Information

#### Supplemental Text

##### Sex-stratified datasets

Sex-stratified genome-wide association study (GWAS) summary statistics for most traits were based on published data obtained by downloading from public repositories (e.g. http://www.med.unc.edu/pgc/results-and-downloads) or directly from analysts (see **Table 1** for details), with several exceptions, as described below.

PGC data for schizophrenia, major depressive disorder (MDD), and bipolar disorder were processed using a unified quality control pipeline as part of a separate study [Blokland et al, in preparation]. As part of this pipeline, samples were removed if they showed relatedness within or between datasets (PI-HAT <0.1). A subset of the MDD samples with recurrent MDD were also used for a sex-stratified analysis.

Unpublished sex-stratified GWAS summary-statistics for anxiety diagnosis, MDD diagnosis, and current and previous smoking were downloaded from http://www.nealelab.is/uk-biobank. The UK Biobank (UKBB) phenotype codes that were used are: “self-reported anxiety/panic attacks” (20002_1287), “self-reported depression” (20002_1286), “smoking status previous” (20116_1), and “smoking status current” (20116_2). Sex-stratified summary data from 2 other unpublished sex-stratified phenotypes in UK Biobank (neuroticism and cannabis use) were also available from co-authors; the UK Biobank codes that were used are: “neuroticism total score of 12 dichotomous items of the Eysenck Personality Questionnaire Revised Short Form” (20127) and "ever taken cannabis” (20453).

For ASD, 2 datasets were meta-analysed based on data from 2 publications (Mitra et al. 2016; Grove et al. 2019) using an inverse-variance weighted fixed effects model, implemented in METAL (Willer, Li, and Abecasis 2010); the genetic correlation (in LDSC (Bulik-Sullivan et al. 2015)) between these 2 ASD datasets was: rg(SE)=0.70(0.11) in males and could not be estimated in females due to low sample size.

For 2 phenotypes, data were available from 2 different sources and we assessed the genetic correlations between them and used only one of the datasets going forward; for MDD, data were available from the PGC and the UK Biobank (LDSC rg(SE)=1.16(0.37) for males & 0.60(0.11) for females) and we selected the PGC data as it was based on a rigorously phenotyped clinical case-control dataset; for alcohol consumption data were available from 2 published studies (Clarke et al. 2017; Schumann et al. 2016) (rg(SE)=0.66(0.11) for males & 1.18(0.27) for females) and we used data from the larger study (Clarke et al. 2017).

**Quality control & processing datasets for analyses**

The majority of available datasets had already undergone rigorous quality control, as described in each study publication (see **Table 1** for references). We performed several additional steps, as follows. All datasets were on the same genomic build (hg19). SNPs were filtered for minor allele frequency (MAF), imputation quality, and sample size (N.B. different filters were applied for LDSC & meta-analyses; see below).

*LDSC*

The default parameters of LDSC (as per the script munge_sumstats.py) were used based on available columns provided in the shared datasets, removing variants as follows: MAF<=0.01, INFO<=0.9, N<(90th percentile N)/1.5, indels, strand-ambiguous SNPs and variants not in HapMap-3. Where the sample size was not provided per variant, the total sample size was provided to LDSC.

**References**


Blokland, GAM, Grove, J, Chen, CH, Cotsapas, C, Tobet, S, Handa, R, Schizophrenia working group of the Psychiatric Genomics Consortium, Bipolar Disorder working group of the Psychiatric Genomics Consortium, Major Depressive Disorder working group of the Psychiatric Genomics Consortium, Lundbeck Foundation Initiative for Integrative Psychiatric Research (iPSYCH), Damm Als T, Anders D. Børglum, AD, Smoller, JW, Petryshen, TL, Goldstein, JM. Sex-Dependent Shared and Non-Shared Genetic Architecture Across Mood and Psychotic Disorders. In Preparation.

Bulik-Sullivan, Brendan K., Po-Ru Loh, Hilary K. Finucane, Stephan Ripke, Jian Yang, Schizophrenia Working Group of the Psychiatric Genomics Consortium, Nick Patterson, Mark J. Daly, Alkes L. Price, and Benjamin M. Neale. 2015. “LD Score Regression Distinguishes Confounding from Polygenicity in Genome-Wide Association Studies.” Nature Genetics 47 (3): 291–95.

Clarke, T-K, M. J. Adams, G. Davies, D. M. Howard, L. S. Hall, S. Padmanabhan, A. D. Murray, et al. 2017. “Genome-Wide Association Study of Alcohol Consumption and Genetic Overlap with Other Health-Related Traits in UK Biobank (N=112 117).” Molecular Psychiatry 22 (10): 1376–84.

Grove, Jakob, Stephan Ripke, Thomas D. Als, Manuel Mattheisen, Raymond K. Walters, Hyejung Won, Jonatan Pallesen, et al. 2019. “Identification of Common Genetic Risk Variants for Autism Spectrum Disorder.” Nature Genetics 51 (3): 431–44.

Mitra, Ileena, Kathryn Tsang, Christine Ladd-Acosta, Lisa A. Croen, Kimberly A. Aldinger, Robert L. Hendren, Michela Traglia, et al. 2016. “Pleiotropic Mechanisms Indicated for Sex Differences in Autism.” PLoS Genetics 12 (11): e1006425.

Schumann, Gunter, Chunyu Liu, Paul O’Reilly, He Gao, Parkyong Song, Bing Xu, Barbara

Ruggeri, et al. 2016. “KLB Is Associated with Alcohol Drinking, and Its Gene Product β-Klotho Is Necessary for FGF21 Regulation of Alcohol Preference.” Proceedings of the National Academy of Sciences, November. https://doi.org/10.1073/pnas.1611243113.

Willer, Cristen J., Yun Li, and Gonçalo R. Abecasis. 2010. “METAL: Fast and Efficient Meta-Analysis of Genomewide Association Scans.” Bioinformatics 26 (17): 2190–91.

#### Supplemental Table Titles

Table S1: Data availability.

Table S2: Sex-specific population prevalence estimates from USA & Denmark.

Table S3: ICD codes used to estimate sex-specific population prevalence in USA.

Table S4: Observed & liability scale SNP-based h2 estimated by sex using LDSC.

Table S5: Observed & liability scale SNP-based h2 estimated by sex using LDAK.

Table S6: Genetic correlation estimates (LDSC) across sex for each pair of traits.

Table S7: Results of top genes with sex-differentiated effects.

Table S8: Results of gene set enrichment analysis.

Table S9: Genetic correlation estimates (LDSC) within sex, across each pair of traits.

#### Supplemental Figures

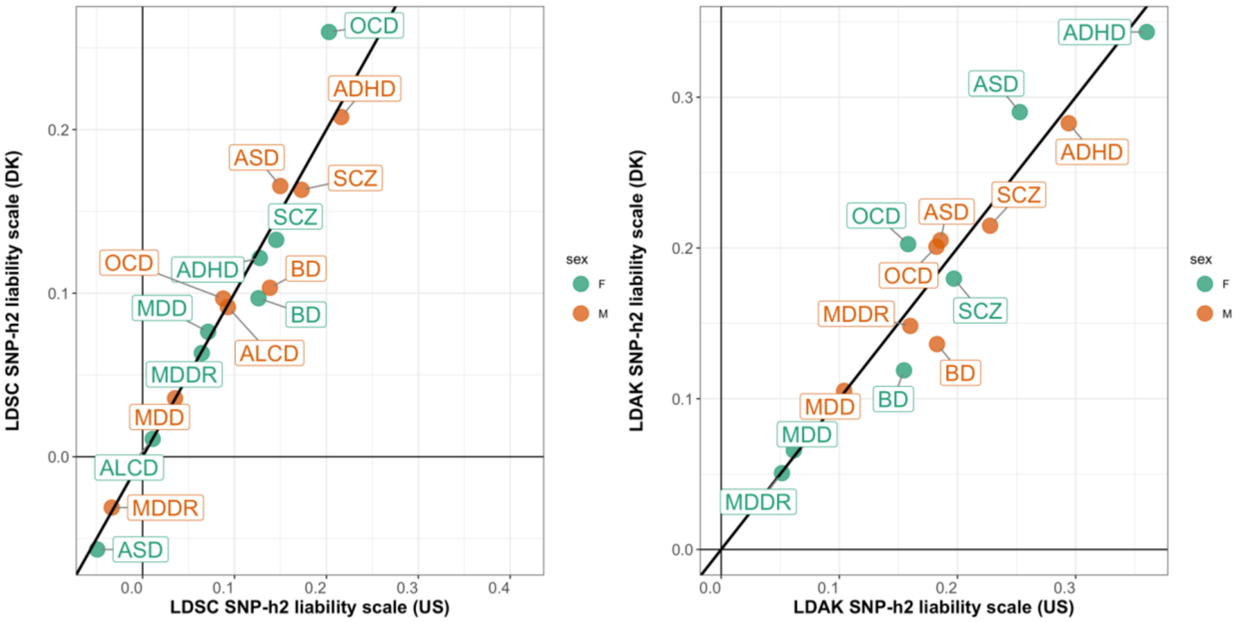


**Figure S1.** Scatter plots of the correlation of SNP-based $h^2$ estimates using US- and Denmark- (DK) based population prevalence rates, for 2 different SNP-based $h^2$ estimation methods: (A) LDSC and (B) LDAK. Results show high correlation when using US-based (x-axis) and DK-based (y-axis) prevalence estimates for LDSC ($r^2$=0.97, p=5.1x$10^{-10}$) and LDAK ($r^2$=0.95, p=1.2x$10^{-8}$). Female SNP-based $h^2$ estimates are shown in green and male SNP-based $h^2$ estimates are shown in orange. The X:Y line is plotted in black. Standard errors (SE) are reported in the Tables S3 & S4. Phenotype abbreviations are listed in Table 1.

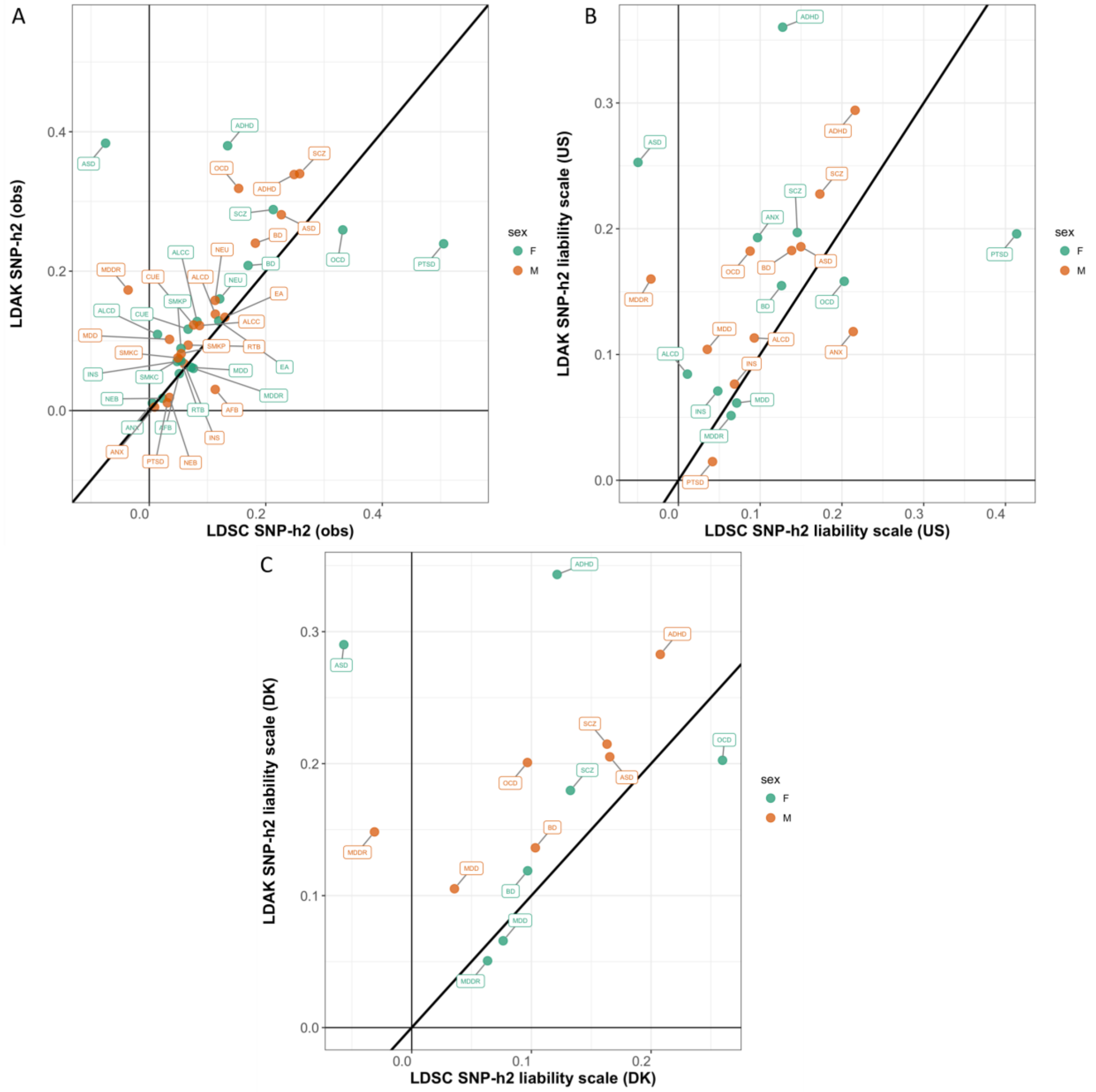


**Figure S2.** Correlation of SNP-based $h_2$ estimates using LDSC and LDAK methods. Scatter plots of SNP-based $h_2$ estimates on the (A) observed scale, (B) liability scale using US-based prevalence, and (C) liability scale using DK-based prevalence; each using LDSC (x-axis) and LDAK (y-axis). The X:Y line is plotted in black. Observed scale SNP-based $h_2$ estimates from LDAK and LDSC methods are modestly correlated: $r_2$=0.53, p=4.4x10$_{-4}$ for all traits, $r_2$=0.73, p=4.1x10$_{-7}$ excluding traits for which SNP-based $h_2$ could not be reliably estimated in LDSC, i.e. PTSD & MDDR in males and ASD & ALCD in females. The liability scale SNP-based $h_2$ estimates are only modestly correlated; US: $r_2$=0.32, p=0.15 for all traits, $r_2$=0.42, p=0.08 excluding the 4 traits; DK: $r_2$=0.22, p=0.44 for all traits, $r_2$=0.60, p=0.04 excluding the 4 traits. Standard errors (SE) are reported in the Tables S3 & S4. Phenotype abbreviations are listed in Table 1.

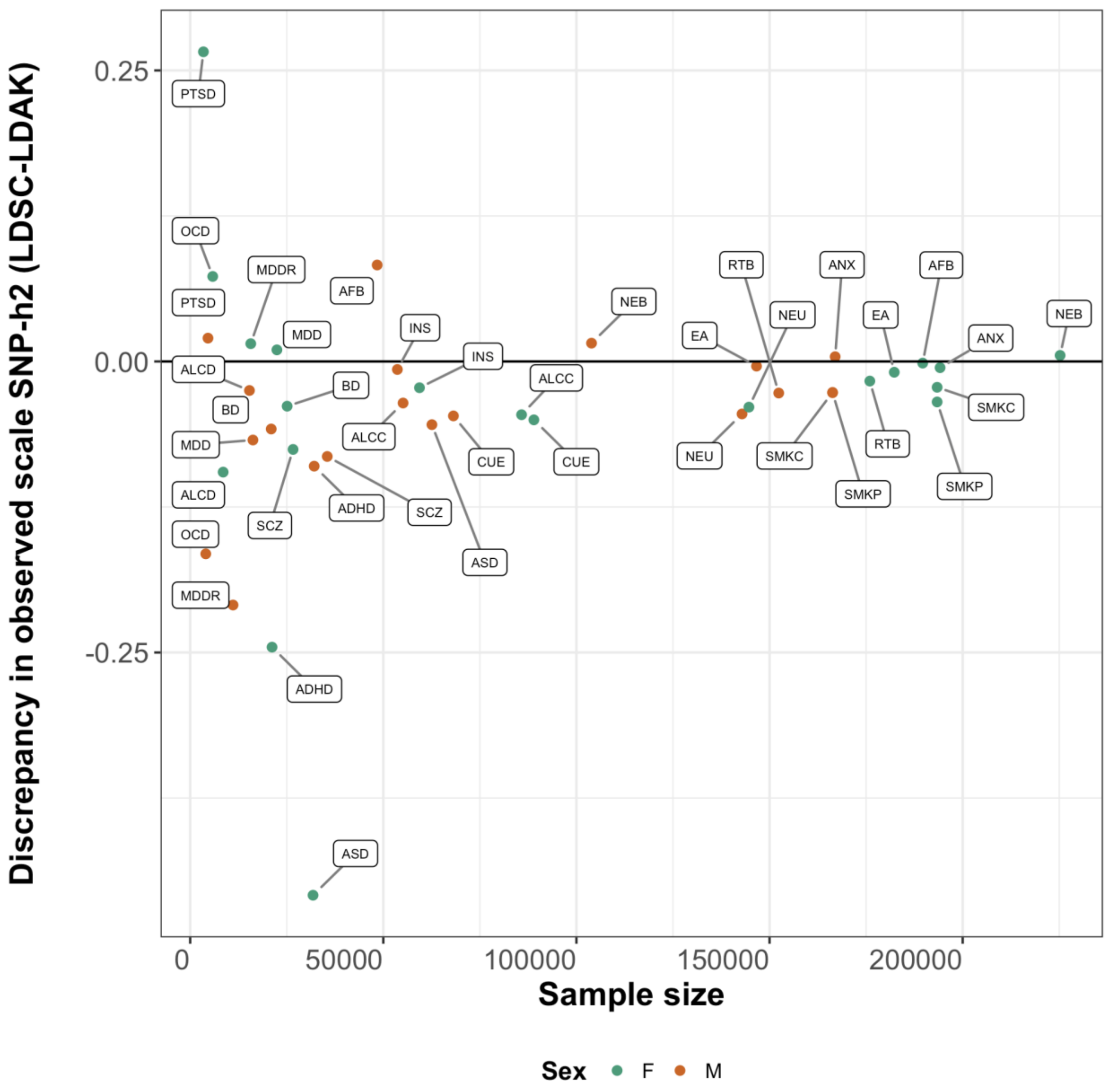


**Figure S3.** The effect of sample size on the difference between the LDSC and LDAK methods. Each point represents a value for the difference in SNP-based $h_2$ estimated using two methods (LDSC and LDAK) for females (green) and males (orange). The sample size of the cohorts is shown on the x-axis, and the difference between SNP-based $h_2$ estimates on the observed scale is shown on the y-axis. Phenotype abbreviations are listed in Table 1.

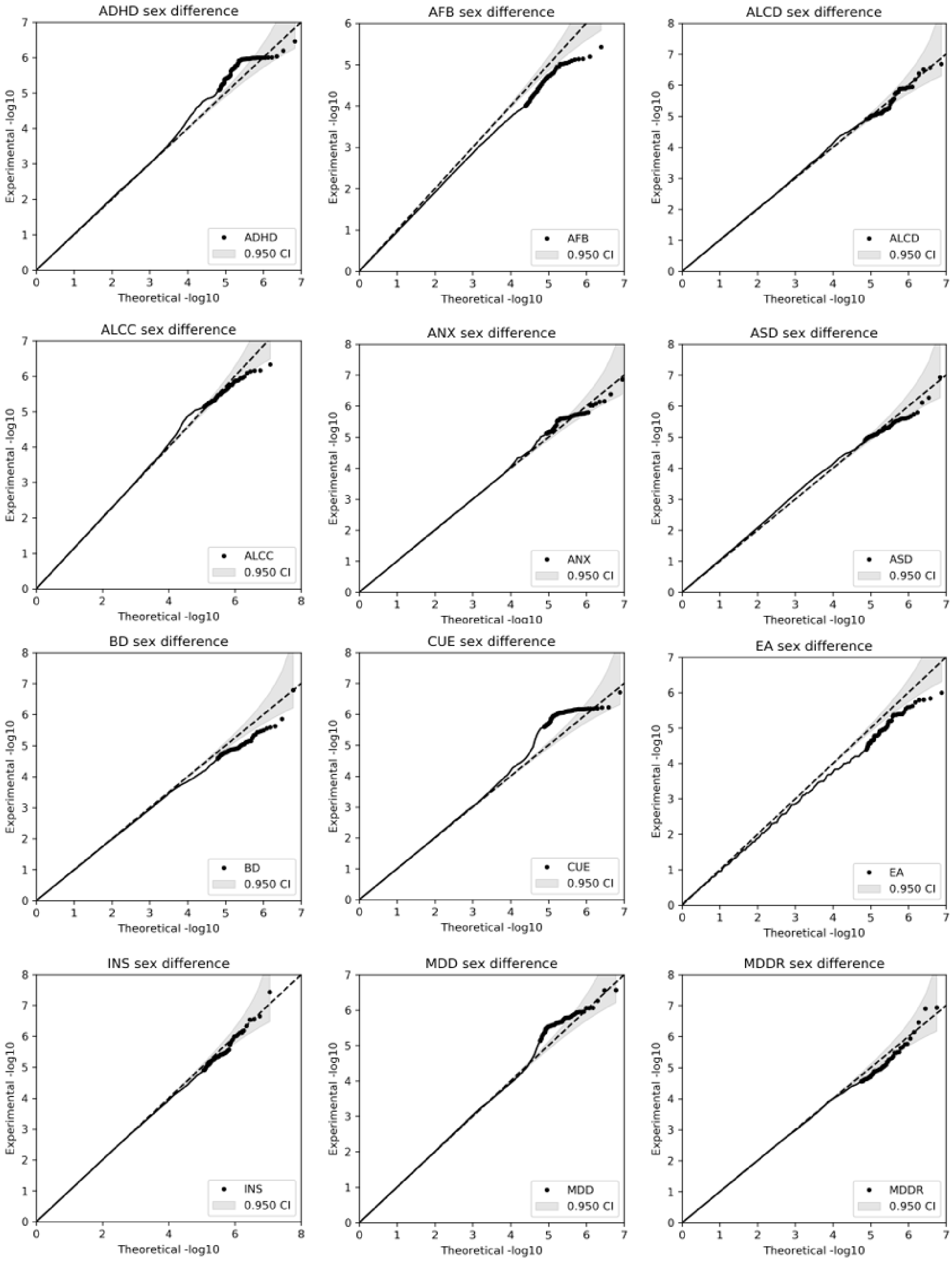

ADHD sex difference
AFB sex difference
ALCD sex difference
ALCC sex difference
ANX sex difference
ASD sex difference
BD sex difference
CUE sex difference
EA sex difference
INS sex difference
MDD sex difference
MDDR sex difference
Experimental -log10
Theoretical -log10
ADHD
AFB
ALCD
ALCC
ANX
ASD
BD
CUE
EA
INS
MDD
MDDR
0.950 CI

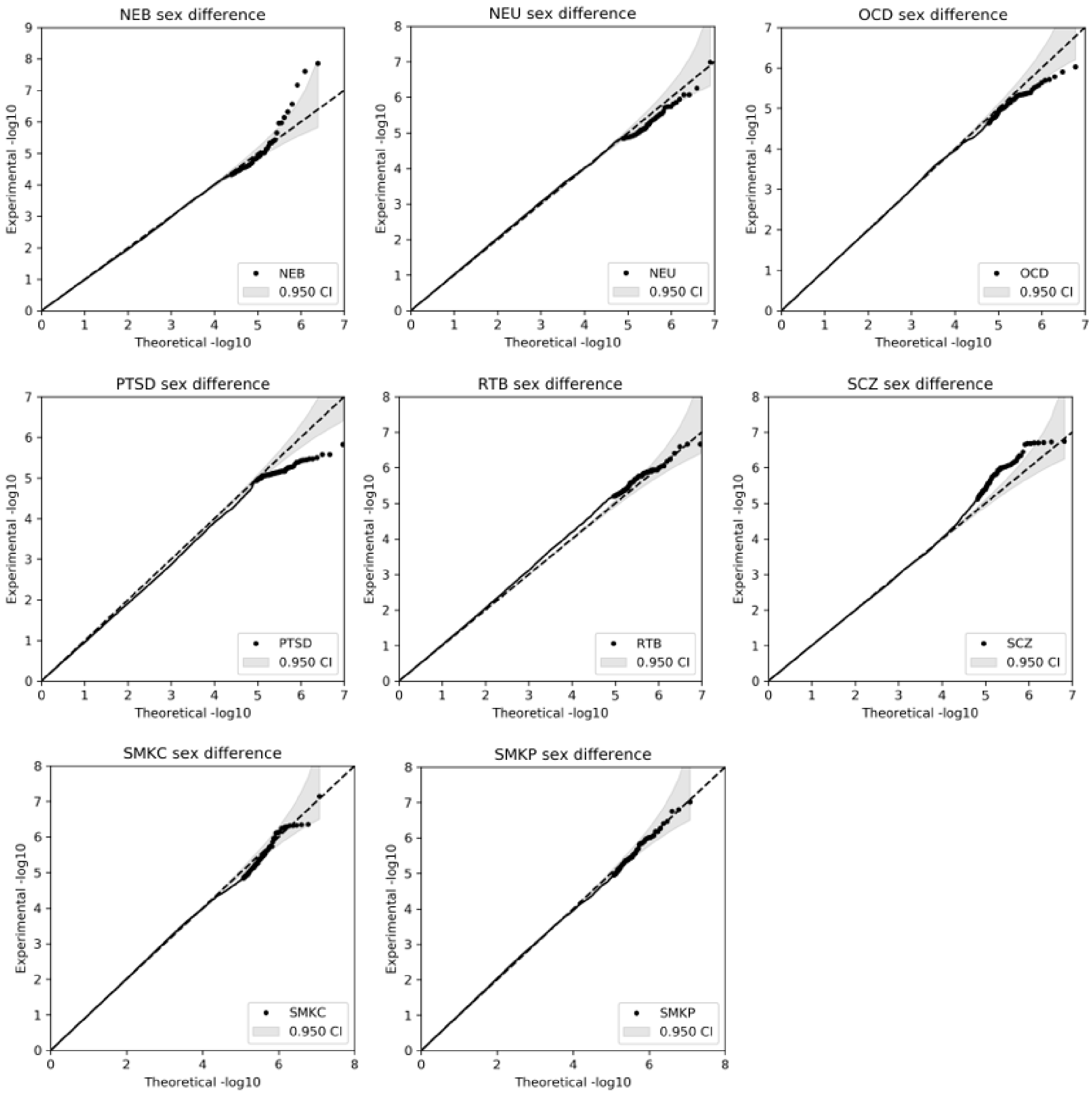


**Figure S4.** Quantile-quantile (QQ) plots of z-score p-values. Theoretical -log10(p-values) are shown on the x-axis, and experimental -log10(p-values) are shown on the y-axis. The X:Y line is plotted as a black dashed line and the grey area represents the 95% confidence interval. Phenotype abbreviations are listed in Table 1.

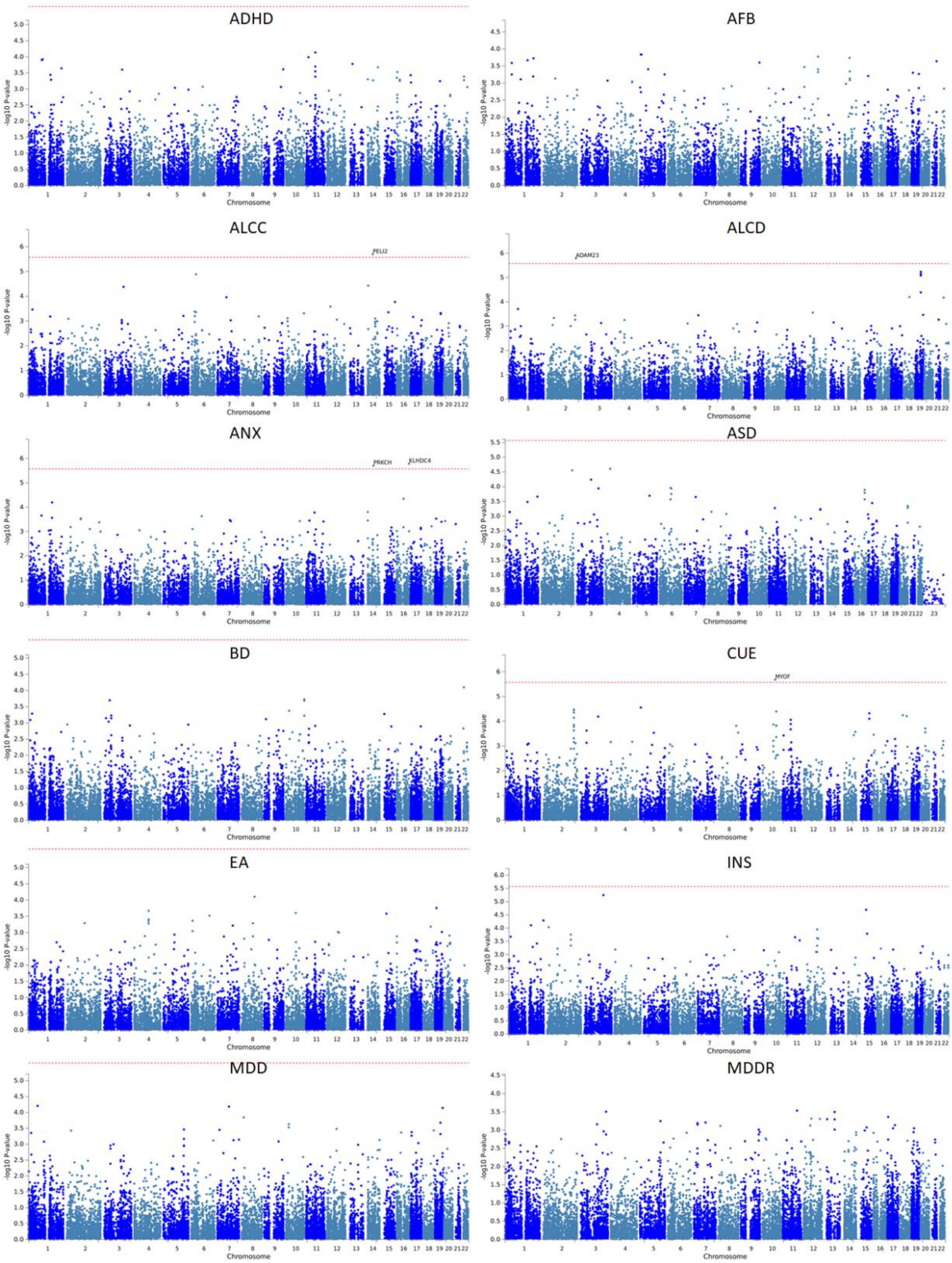
ADHD
AFB
ALCC
PELI2
ALCD
ADAM23
ANX
PRKCH
KLHDC4
ASD
BD
CUE
MYOF
EA
INS
MDD
MDDR
-log10 P-value
Chromosome

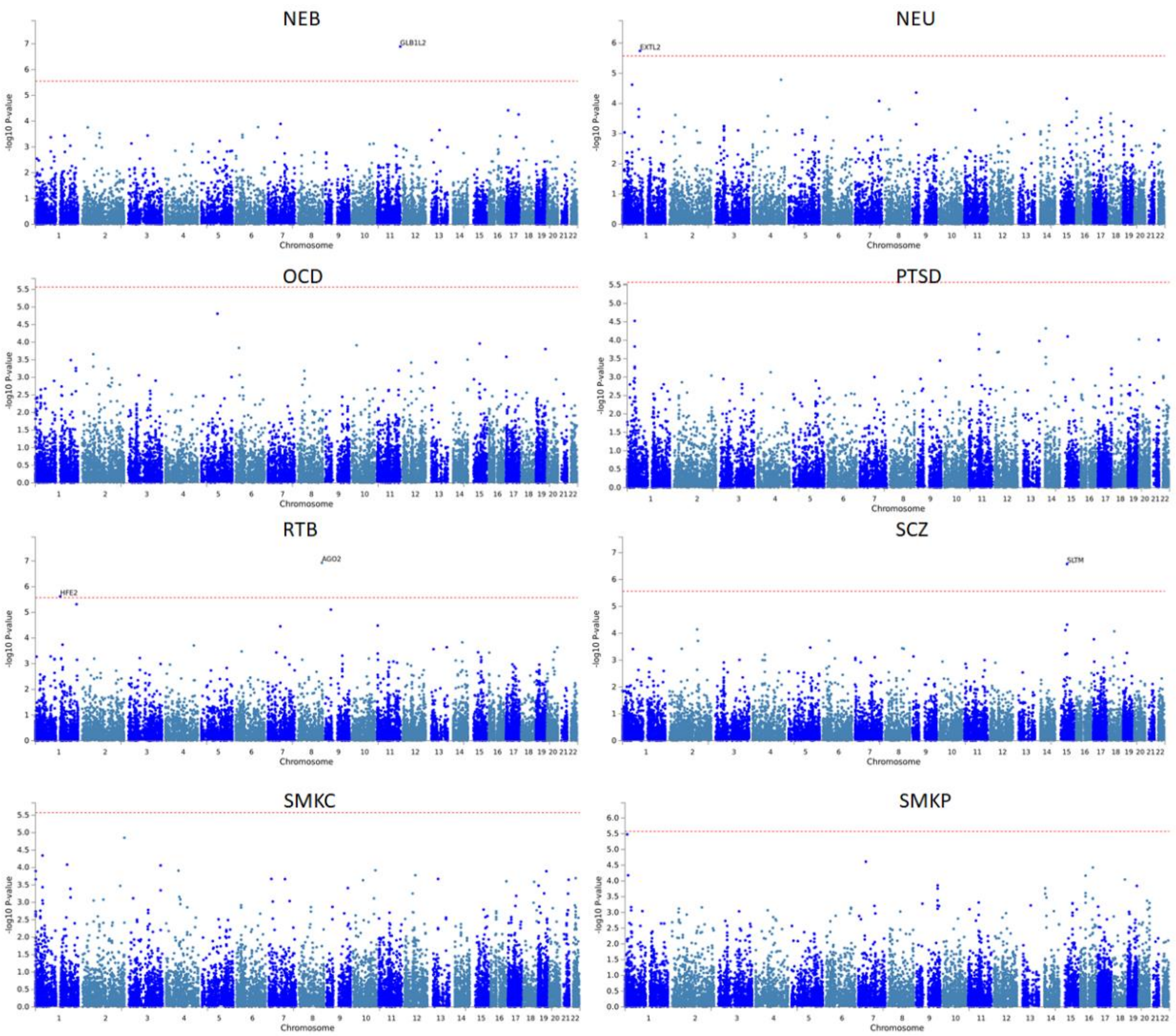


**Figure S5.** Manhattan plots of gene-based analysis in FUMA using z-score p-values. Each point represents a gene. Chromosome and base pair positions are shown on the x-axis, while the -log10(p-values) are shown on the y-axis. Phenotype abbreviations are listed in Table 1.

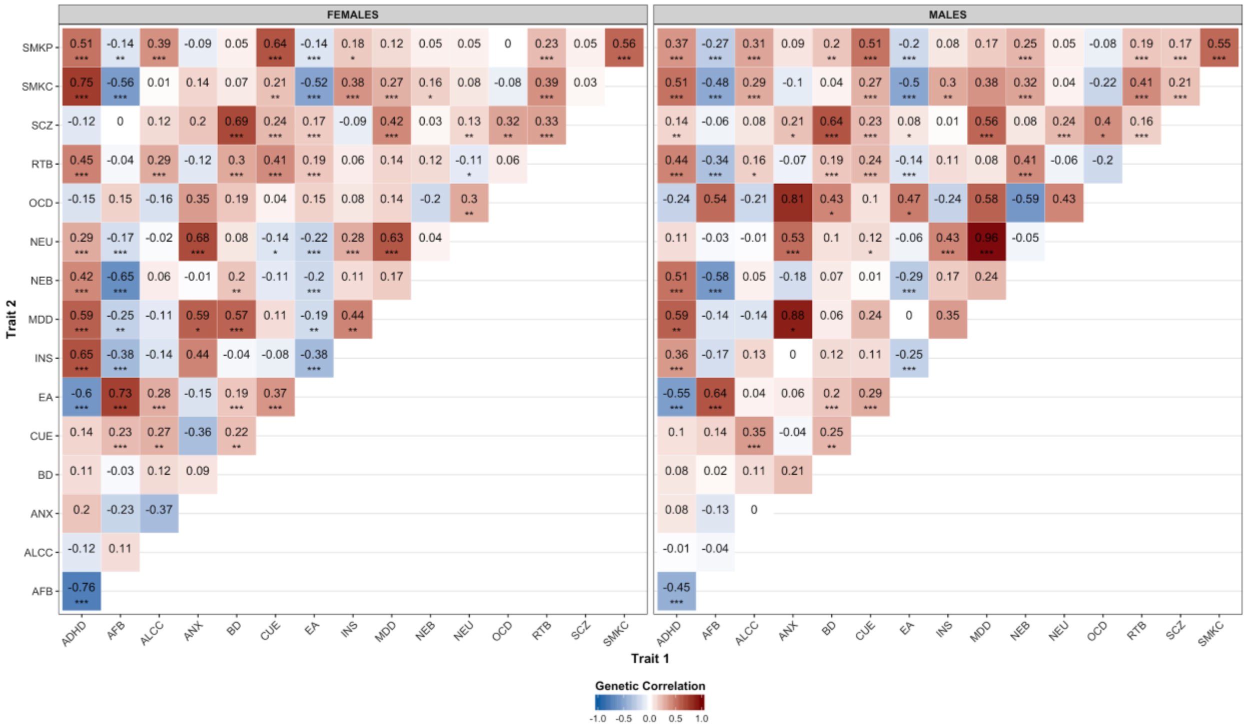


**Figure S6.** Heatmaps of genetic correlation analyses for females (left) and males (right). Red represents positive correlations and blue represents negative correlations. The correlation coefficient is listed in each cell. The significance of the correlations is denoted with asterisks, as follows: * p<0.05, **p<0.01, ***p<0.001. Phenotype abbreviations are listed in Table 1.
